# Starvation Response and RNA Polymerase Guide Antibiotic-induced Mutable Gambler Cells

**DOI:** 10.1101/2022.07.06.499071

**Authors:** Yin Zhai, P.J. Minnick, John P. Pribis, Libertad Garcia-Villada, P.J. Hastings, Christophe Herman, Susan M. Rosenberg

## Abstract

Antibiotics can induce mutations that cause antibiotic resistance, triggered by stress responses. The identities of the stress-response activators reveal environmental cues that elicit mutagenesis, and are weak links in mutagenesis networks, inhibition of which could slow evolution of resistance during antibiotic therapies. Despite pivotal importance, few identities and fewer functions of stress responses in mutagenesis are clear. Here, we identify the stringent starvation response in fluoroquinolone-antibiotic ciprofloxacin (stress)-induced mutagenesis. We show its role in transient differentiation of a mutable “gambler” cell subpopulation. Stringent activity is induced by reactive oxygen and upregulates two small (s)RNAs that induce the general stress response, which allows mutagenic DNA-break repair. Surprisingly, RNA polymerase and response-activator nucleotide (p)ppGpp promote ciprofloxacin-induced DNA-damage signaling. We propose a model for their regulation of DNA-damage processing in transcribed regions. The data demonstrate a critical node in ciprofloxacin-induced mutagenesis, imply RNA-polymerase regulation of repair, and identify promising targets for resistance-resisting drugs.

## INTRODUCTION

Antibiotic resistance is a global health threat with an estimated 1.27 million deaths worldwide resulting from antibiotic resistance in 2019 (Murray et al., 2022). Antibiotic resistance results from uptake of resistance genes from other bacteria or mutation of native genes (Blazquez et al., 2018). New mutations drive resistance to diverse antibiotics and dominate antibiotic resistance among the World Health Organization’s priority pathogens (Govindaraj Vaithinathan and Vanitha, 2018). Many current antimicrobial therapies can induce mutagenesis that leads to resistance (Cirz et al., 2005; Gutierrez et al., 2013; Kohanski et al., 2010), compounding the problem.

Resistance has been circumvented, historically, by development of new antibiotics, and strategies in novel antibiotic development are promising (Martin et al., 2020; Wang et al., 2022). However, the rates of evolution of resistance far exceed rates of introduction of new antibiotic drugs (Ardal et al., 2020; Davies and Davies, 2010). A complementary strategy could prolong the utility of new and old antibiotics: to discover, then inhibit, molecular mechanisms that drive evolution of resistance, e.g., (Al Mamun et al., 2012; Cirz et al., 2005; Rosenberg and Queitsch, 2014). The molecular mechanism of fluoroquinolone antibiotic-induced mutagenesis is unique in being supported by previous identification of a large network of proteins that promote stress-induced mutagenic DNA-break repair (MBR) (Al Mamun et al., 2012), and its induction by the fluoroquinolone antibiotic ciprofloxacin (cipro) (Pribis et al., 2019), reviewed (Pribis et al., 2022; Waylen et al., 2020). Cipro-induced mutagenesis, therefore, provides a useful framework and model system for strategic inhibition of antibiotic-induced mutagenesis, including elucidation of the molecular mechanism and its regulation.

Fluoroquinolones bind and inactivate type-II topoisomerases mid-reaction, which kills cells by inflicting DNA double-strand breaks (DSBs) (Drlica, 1999). Type II topoisomerases relieve replication- and transcription-induced positive DNA supercoils by cleaving opposite DNA strands, passing a duplex through, then ligating the DSB ends (Wang, 1998). Survival of fluoroquinolone-treated bacteria therefore requires DSB-repair proteins and the SOS DNA-damage response (Anderson and Osheroff, 2001), which upregulates repair. Fluoroquinolone resistance occurs mostly by *de novo* mutations that alter the topoisomerase, preventing drug binding, or upregulate efflux pumps (Blair et al., 2015; Cullen et al., 1989; Mehrad et al., 2015; Oethinger et al., 1998; Oram and Fisher, 1991; Ubukata et al., 1989; Zhang et al., 2011). Cipro, the most commonly prescribed fluoroquinolone (Werner et al., 2011), induces the SOS response (Howard et al., 1993) and mutagenesis (Cirz et al., 2005; Pribis et al., 2019) at low, “sub-inhibitory” doses, activating a stress-induced MBR mechanism (Pribis et al., 2019). A detailed pathway of transient differentiation of a mutable “gambler” cell subpopulation is described (Figure 1A) (Pribis et al., 2019), in which cipro induces topoisomerase-mediated DSBs, which induce the SOS response in all cells; SOS induces ubiquinone (electron-transfer)-dependent reactive-oxygen species (ROS) in a roughly 20% cell subpopulation. The ROS induce two small RNAs, DsrA and ArcZ, which activate the general stress response by binding and stabilizing the mRNA encoding its transcriptional activator, sigma S (σ^S^), which licenses MBR (Lombardo et al., 2004). σ^S^ executes a switch from high-fidelity, to mutagenic DSB repair (Ponder et al., 2005; Shee et al., 2011) by allowing SOS-upregulated error-prone DNA polymerases to participate in repair, and leave errors that persist as mutations near DSB-repair sites (Ponder et al., 2005; Shee et al., 2012). The gambler cells “risk” mutagenesis while cells outside the subpopulation remain stable.

**Figure 1.**
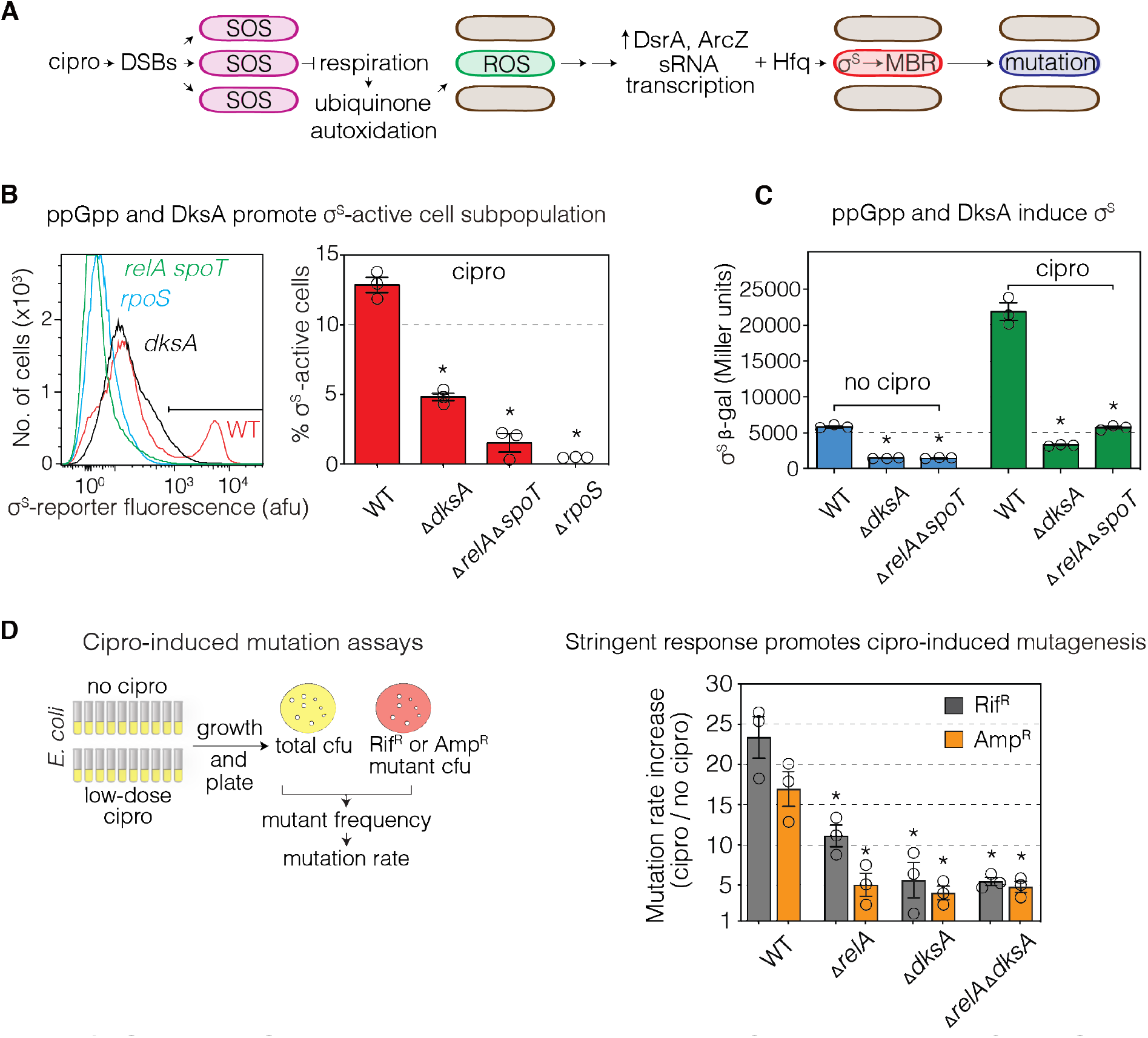
Stringent Starvation-Response Activators in Cipro Induction of the General Stress Response and Mutagenesis. (A) Summary diagram of observations of (Pribis et al., 2019) that cipro induces DSBs and the SOS response in all cells; the SOS response then promotes ROS in a ∼20% cell subpopulation. The ROS induce transcription of two small (s)RNAs, DsrA and ArcZ, which allow translation of *rpoS* mRNA into σ^S^ protein, thereby creating the ∼20% σ^S^- (general stress response)-active “gambler” cell subpopulation that, produces nearly all cipro-induced mutants via mutagenic break repair (MBR) (Pribis et al., 2019). (B) Stringent-response activators (p)ppGpp and DksA are required for genesis of the σ^S^-active cell subpopulation. Data at 24 h MAC cipro, stationary phase. σ^S^-active cells, right of the gate (black bar, Methods). Δ*relA ΔspoT* cells lack (p)ppGpp synthases. (C) Cipro induction of σ^S^ protein requires (p)ppGpp and DksA. β-galactosidase activity, *rpoS-lacZ* translational fusion. Data at 24 h MAC cipro, stationary phase. (D) Cipro-induced mutagenesis promoted by stringent-response activators DksA protein and RelA (p)ppGpp synthase. DksA deletion reduced cipro-induced mutagenesis by 78% ± 8% for RifR, and 77% ± 2% for AmpR. Loss of RelA did so by 52% ± 2% for RifR and 71% ± 5% for AmpR mutagenesis. Means ± SEM, 3 experiments. *p<0.001 (B, C), *p<0.01 (D), one-way ANOVA with Tukey’s post-hoc test.

We report here the identification and function of a second critical regulatory node in the transient differentiation of mutable gambler cells: the bacterial starvation, or stringent, stress response. We document its necessity in cipro-induced MBR, and in formation of gambler cells. Surprisingly, we also discover a role of the stringent response-activating “alarmone,” guanosine tetra- (or penta-) phosphate, (p)ppGpp, and its target RNA polymerase (RNAP) in facilitating cipro-induced DNA-damage signaling that activates the SOS response. The data imply that RNAP and transcription contribute to much of DNA-damage signaling induced by cipro. The data identify a critical regulatory hub, and potential druggable targets in cipro-induced mutagenesis to antibiotic resistance (Cirz et al., 2005) and cross resistance to antibiotics not yet encountered (Pribis et al., 2019), and a novel role of stringent-response activation in DNA-damage signaling. Non-redundant regulatory hubs are candidate targets for drugs to slow evolution of resistance, which become accessible as molecular mechanisms are uncovered.

## RESULTS

### Starvation-response Activators Guide Pathway to σ^S^–active Gambler Cells

The stringent response transcriptionally upregulates hundreds of genes among which are *rpoS*, encoding σ^S^ protein, DsrA sRNA, and mRNAs of proteins that prevent σ^S^ degradation (Battesti et al., 2015; Girard et al., 2018). Some but not all inducers of the σ^S^ response are signaled via the stringent response. Data in Figure 1B show that stringent-response activators DksA protein and (p)ppGpp promote cipro induction of the σ^S^-activated gambler cell subpopulation. RelA and SpoT are (p)ppGpp synthases, so deletion of their genes prevents synthesis of the phosphorylated nucleoside. The σ^S^-active cell subpopulation (Figure 1B), and bulk levels of σ^S^-protein (Figure 1C), are decreased sharply in cells that lack the stringent-response activators. We conclude that the stringent response is required for cipro induction of σ^S^.

Using fluctuation-test assays (Figure 1D, left), we found that the stringent response promoted cipro-induced mutagenesis (Figure 1D, right), producing rifampicin- or ampicillin-cross-resistant mutants (RifR or AmpR), per (Pribis et al., 2019). These are caused by specific base substitutions in the *rpoB* gene, or any null mutation in *ampD*, respectively. DksA and RelA, i.e., (p)ppGpp, were also required for half to over two thirds of cipro-induced RifR and AmpR mutagenesis (Figure 1D and legend), demonstrating a stringent-response role in cipro-induced MBR.

### A ROS-induced Stringent Response-on Cell Subpopulation Activates sRNA Transcription

We created a fluorescence transcriptional reporter (Minnick, 2019) of the (p)ppGpp-activated gene *rmf* (ribosome modulation factor) (Izutsu et al., 2001), which allows quantification of stringent-response activity in single cells. In Figure 2A, “sub-inhibitory” cipro at minimum antibiotic concentration (MAC, 10% survival, Materials and Methods) induced the stringent response in a 16% ± 2% subpopulation of cells, similar in size to the ROS-high and σ^S^–active gambler-cell subpopulations induced by cipro (Pribis et al., 2019). ROS were required for the cipro-induced stringent response-active cell subpopulation, which was reduced by addition of the ROS-quenching agent thiourea (Figure 2B, TU), which scavenges hydroxyl radical (Novogrodsky et al., 1982; Touati et al., 1995). We also found that DksA and (p)ppGpp synthases were required for cipro- (and ROS-)induced transcription (Pribis et al., 2019) of the σ^S^-activating sRNAs DsrA and ArcZ (Figure 2C). The two sRNAs did not affect induction of the stringent response by ROS (Figure 2D). We conclude that cipro-induced ROS induce the stringent response, which then promotes synthesis of the sRNAs required for the σ^S^-active cell subpopulation (summarized, Figure 2E).

**Figure 2.**
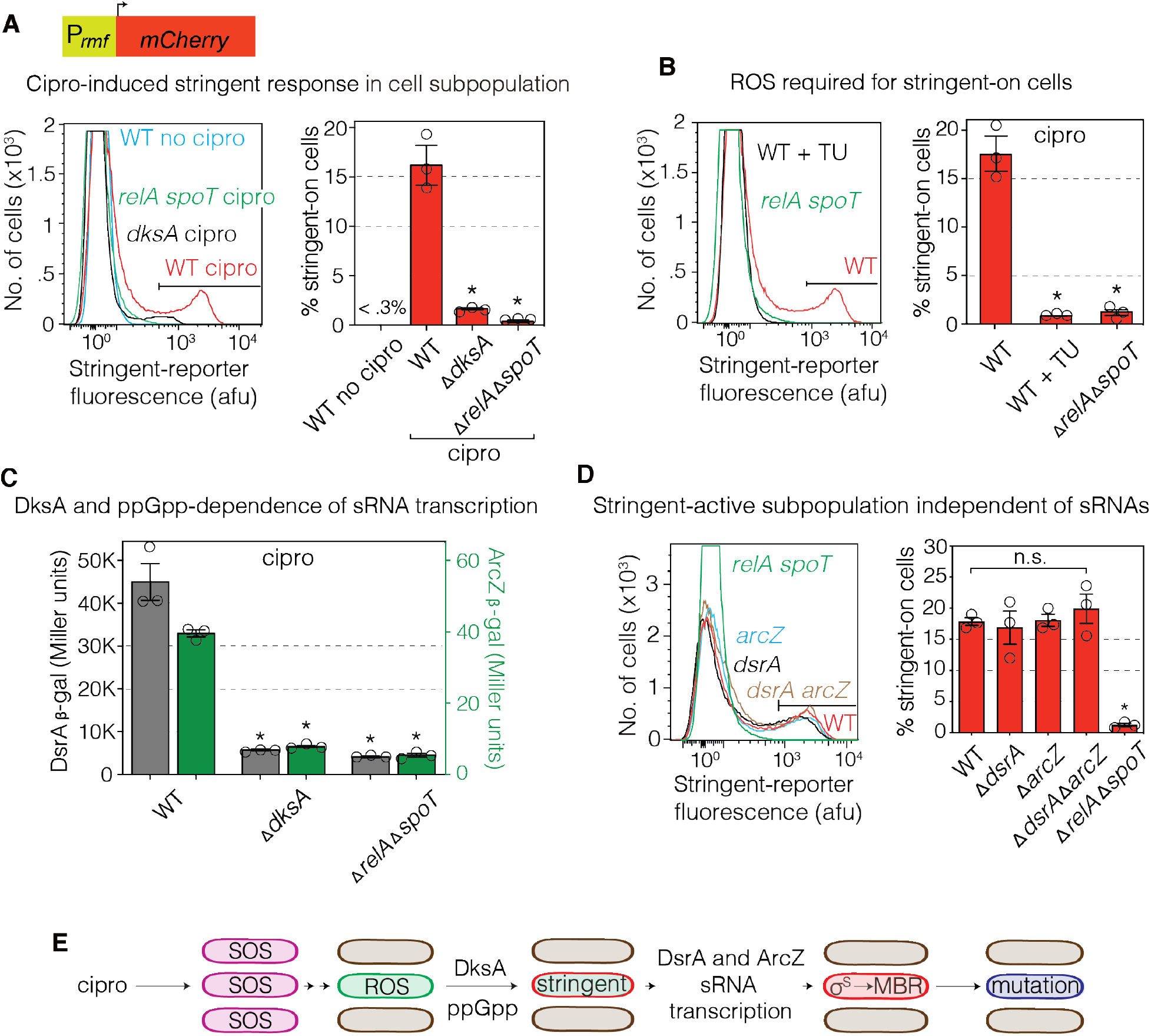
Cipro-induced Stringent Response Occurs in a Cell Subpopulation. (A) Upper: diagram of stringent-response fluorescence-reporter gene engineered into chromosomal *att*λ site (Minnick, 2019). Lower: Cipro induces stringent-response activity in a cell subpopulation. Data at 16 h MAC cipro, log phase. (B) ROS are required for the cipro-induced stringent response. ROS scavenger thiourea (TU) reduces the stringent response-on cell subpopulation. Data at 16 h MAC cipro, log phase. (C) Transcriptional induction of sRNA-gene *dsrA* and *arcZ* promoters requires DksA protein and (p)ppGpp. MAC cipro, at 12-14 h, log phase. β-galactosidase activity, P_*dsrA*_*lacZ* and P_*arcZ*_*lacZ* reporters. (D) σ^S^-activating sRNAs DsrA and ArcZ are induced by the stringent response (C) and are not required for stringent-response induction. Data at 16 h MAC cipro, log phase. That is, the stringent response acts upstream of induction of these sRNAs in the MBR pathway. (A, B, D) Flow cytometry of cells carrying stringent-response reporter P_*rmf*_*mCherry*. Stringent response-on cells, right of the gate (black bar, Methods). Representative histograms and means ± SEM, 3 experiments, right. *p<0.001 (A-D), one-way ANOVA with Tukey’s post-hoc test; n.s. not significant. (E) Summary: The stringent response is activated by cipro-induced ROS in a cell subpopulation and induces the sRNAs required for formation of σ^S^-active gamblers and MBR mutagenesis.

### ROS-high Cells Become Stringent-induced

We found that the ROS-high cells become the stringent response-active cell subpopulation (Figure 3A). ROS and stringent transcriptional activity were followed simultaneously by flow cytometry in time-course experiments in living cells (Figure 3A) via an oxidative stress-response promoter driving GFP, P_*oxyR*_*gfp* (Zaslaver et al., 2006), to infer the presence of ROS in living cells that also carry the stringent-response reporter P_*rmf*_*mCherry*. In Figure 3A, the horizontal flow-cytometry “gates” indicate the gate above which cells are designated ROS-high. The vertical gates divide stringent response-active (or “on”) cells, to the right, from inactive cells (left; Methods).

**Figure 3.**
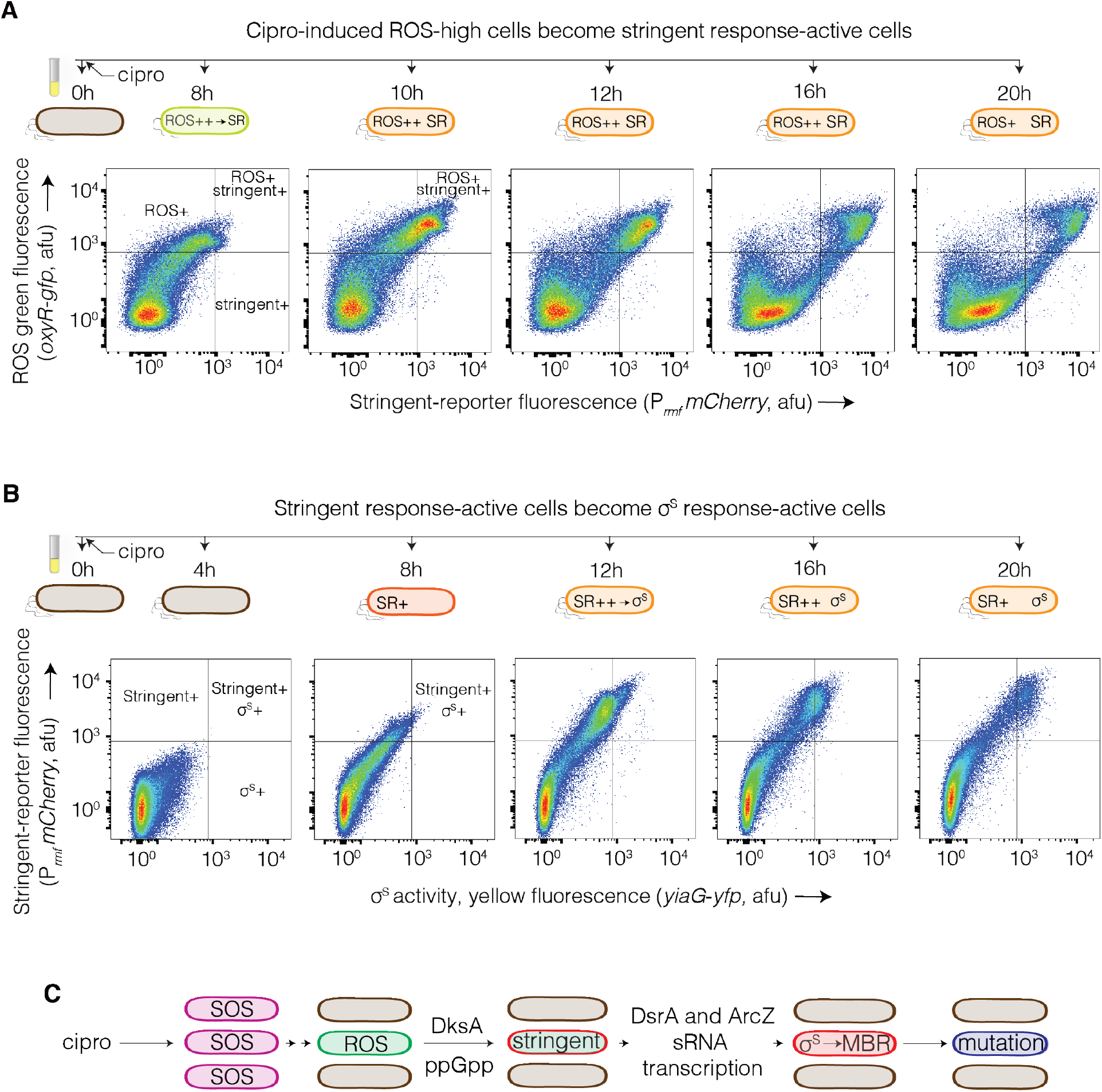
ROS-high Cells Become Stringent-active then σ^S^*-*active Gambler Cells. Time course of stress-response activation in cells with exposure to MAC cipro. (A) Most stringent-response (SR) active cells arise from ROS-high cells. Cipro induction of ROS-high cells, measured with the P_*oxyR*_*gfp* oxidative stress-response reporter, occurs before activation of the stringent response. Then stringent-response activity appears in cells with high ROS: double-positive cells. Eventually some of the stringent-response active cells finally lose ROS, e.g., 16 h and 20 h post addition of cipro to the log-phase cultures. Representative of 3 experiments. (B) Cipro-induced σ^S^ activity arises in stringent-response-on cells. Flow cytometry time course of cells with MAC cipro. Stringent response-on cells precede σ^S^-on cells, and most σ^S^-on yellow cells begin cells as stringent response-on double-positive cells. Double positives, upper right quadrants. Representative of 3 experiments. (C) Summary: There is a transient differentiation of a cipro-induced subpopulation of cells from ROS-high to stringent response-on, to σ^S^-on.

We found that ROS-high cells appeared before stringent activation (Figure 3A), see 8 h after transfer to MAC cipro. Then the ROS-high subpopulation cells became stringent response-active, double-positive cells with few or no stringent-active cells observed that lacked ROS (Figure 3A, 8 - 10 h). The ROS and stringent response were transient. From 12 – 20 h, some of the stringent-active cells appeared to lose high ROS, then stringent-active cells began to decline at 16 h and 20 h (Figure 3A). The transformation from ROS-high to stringent-active double-positive cells occurred progressively (Figure 3A). The data imply that cipro-induced ROS induce the stringent response (Figure 2B) in the same cell subpopulation (Figure 3A).

### Stringent Response-active Cells Become σ^S^*-*active Gamblers

MAC cipro induced the stringent response in a 17% ± 3% cell subpopulation (Figures 2A and 2B, mean ± SEM), similar in size to the σ^S^-active subpopulation (Pribis et al., 2019) (Figure 1B). The following data show that the same cells activate each response, with stringent-on cells becoming σ^S^-active double-positive cells. We moved the *yiaG*-*yfp* chromosomal σ^S^-reporter gene into a strain carrying the chromosomal stringent-response reporter P_*rmf*_*mCherry* and measured cipro-induction of the two stress responses simultaneously (Figure 3B). The first stringent response-active cells were visible at 8 h growth with cipro. Stringent response-active σ^S^-active double-positive cells appeared between 8 h and 20 h, with maximal σ^S^ activity between 12 h and 16 h. At all-time points, nearly all σ^S^-induced cells were stringent response-active, but not all stringent response-active cells were σ^S^-induced (Figure 3B). The data imply that the σ^S^-induced cells, which have been shown to be mutable gamblers (Pribis et al., 2019), arose from a stringent-active cell subpopulation (Figure 3C).

### (p)ppGpp in DNA-damage signaling

Surprisingly, we found that (p)ppGpp, but not DksA, was needed for full cipro-induction of the SOS response (Figure 4A, B, Discussion), and so also for SOS-dependent induction of ROS-high cells (Figure 4C). The data separate the role(s) of DksA and (p)ppGpp in cipro-induced MBR, with DksA functioning exclusively downstream of ROS inducing the sRNAs (Figures 2C, 3C and 4B-D), and (p)ppGpp acting early, promoting SOS induction (summarized Figure 4E). We tested whether the ROS serve any function in MBR other than inducing the stringent response (Figures 2B and 3A) by quenching ROS with TU in Δ*dksA* cells. Mutagenesis is reduced in Δ*dksA* cells and removal of the ROS with TU caused no further reduction (Figure 4D), implying that ROS play no role in cipro-induced mutagenesis other than inducing stringent-response activation, which is DksA dependent, as summarized in Figure 4E.

**Figure 4.**
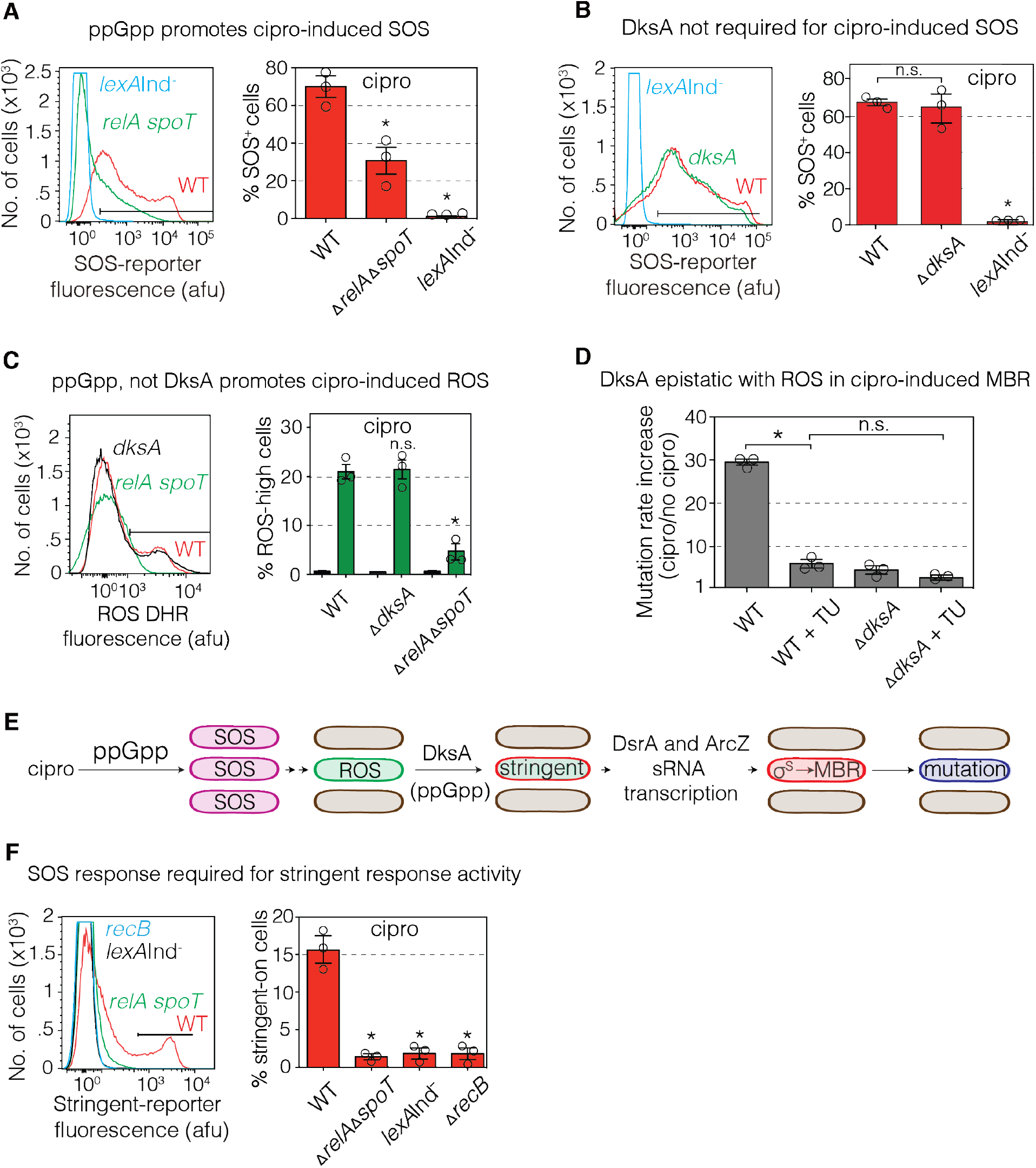
(p)ppGpp and DksA at Separate Steps in Cipro-induced MBR Pathway. (A) (p)ppGpp promotes the cipro-induced SOS response. Flow-cytometry 16 h MAC cipro (log phase). SOS reporter P_*sulA*_*mCherry*. SOS-positive cells, right of the gate (black bar, Methods). (B) DksA-independence of the cipro-induced SOS response. Flow-cytometry 16 h MAC cipro (log phase). SOS reporter P_*sulA*_*mCherry*. SOS-positive cells, right of the gate. (C) Cipro-induced ROS-high cells form (p)ppGpp-dependently and independently of DksA. Flow-cytometry 16 h MAC cipro (log phase). DHR, ROS stain. Green bars stained, black bars not stained. (D) Reduction of ROS causes no further decrease in cipro-induced mutagenesis than is seen in Δ*dksA* cells, i.e., they are epistatic, acting in the same pathway. Thiourea (TU) quenches ROS and most cipro-induced mutagenesis (Pribis et al., 2019) and does not reduce mutagenesis in Δ*dksA* cells further, implying that ROS play no role in the MBR pathway in addition to activation of the stringent response. (E) Summary: the DksA and (p)ppGpp-mediated stringent response is required for cipro-induced σ^S^ response activity and mutagenesis. The stringent response is activated by ROS in the ROS-high cell subpopulation, and induces the sRNAs required for subpopulation cells to become σ^S^-active gamblers, which engage in MBR mutagenesis. (p)ppGpp functions in SOS induction by cipro. (F) Cipro induction of the stringent-response reporter requires SOS induction. Stringent-response-active cells are absent in SOS-uninducible *lexA*Ind^-^ or Δ*recB* cells in log phase growth (16 h). RecBCD enzyme is required for SOS induction by DSBs (McPartland et al., 1980) including those made by cipro interaction with type-II topoisomerases (Pribis et al., 2019). MAC cipro. Flow cytometry, stringent-response reporter P_*rmf*_*mCherry*. Stringent response-active cells, right of the gate (black bar, Methods). Means ± SEM, 3 experiments. *p<0.01 (A), *p<0.001 (B-E), one-way ANOVA with Tukey’s post-hoc test; n.s. not significant.

SOS was required for appearance of the stringent response-active subpopulation (Figure 4F), which fails to form in SOS-deficient *lexA*Ind^-^ or Δ*recB* cells. Our data show that DksA, which promotes the stringent-response transcriptional program, functions after ROS (Figures 3 and 4B, C), in promoting induction of the sRNAs (Figure 2C) required for the appearance of the σ^S^-active cell subpopulation (Pribis et al., 2019). The pathway is summarized in Figure 4E. Because the stringent-response transcriptional program usually uses (p)ppGpp in addition to DksA (Ross et al., 2016; Sanchez-Vazquez et al., 2019), we also indicate (p)ppGpp in parentheses in the DksA-dependent step from ROS to sRNA induction (Figure 4E).

### RNA Polymerase in DNA-Damage Signaling

(p)ppGpp binds two sites in RNA polymerase (RNAP) (Ross et al., 2016; Ross et al., 2013), diagrammed in Figure 5A. Occupancy of site 2 by (p)ppGpp, along with DksA protein binding to RNAP (Figure 5A), allows activation of the stringent-response transcriptional program (Ross et al., 2016; Ross et al., 2013). Occupancy of site 1 by (p)ppGpp stabilizes “backtracked” RNAP (after RNA-error correction) (Sivaramakrishnan et al., 2017). RNAP corrects transcription errors by “backtracking” from the RNA 3’ end, allowing the RNAP nuclease to remove the incorrectly inserted base (Komissarova and Kashlev, 1997). This function is independent of indirect effects from activation of the stringent-response transcriptional program (Kamarthapu et al., 2016). The following data confirm the roles of RNAP in cipro-induced MBR that were implied by the steps promoted by (p)ppGpp in Figures 1-4, above. First, data in Figure 5B show that both sites promote cipro-induced MBR, supporting the hypothesis that the (p)ppGpp (and DksA) mutagenic functions (Figures 1 and 2) operate *via* modulation of RNAP, rather than some other, unknown target of these molecules.

**Figure 5.**
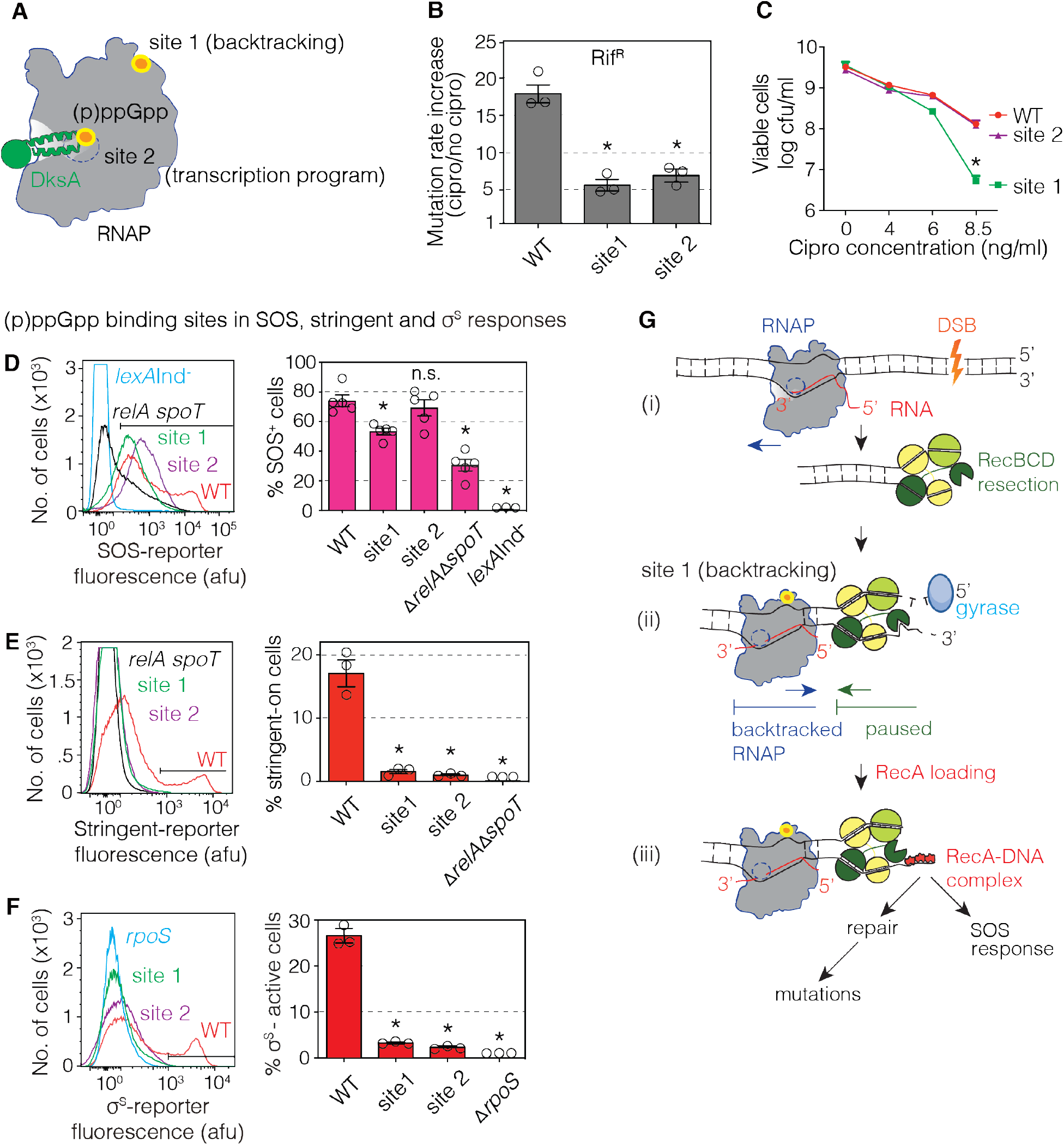
Both (p)ppGpp Binding Sites in RNAP Promote Cipro-induced MBR Pathway. (A) RNA polymerase (RNAP) (p)ppGpp binding sites 1 and 2. (p)ppGpp binding to site 1 stabilizes backtracked RNAP (after RNA-error correction) (Kamarthapu et al., 2016; Ross et al., 2013; Sivaramakrishnan et al., 2017). Site 2 forms with DksA bound to the RNAP secondary channel. RNAP with DksA and (p)ppGpp in site 2 promotes the stringent-response transcriptional program (Ross et al., 2016). (B) Site 1 and site 2 promote cipro-induced mutagenesis. (C) Mutation of RNAP site 1, and not site 2, reduces survival with MAC cipro. 24 h growth in cipro at the concentrations shown. 8.5 ng/ml is the MAC (10% survival) for WT and the site-2 mutant. Means ± range of 2 independent experiments. *p<0.01, one-way ANOVA with Tukey’s post-hoc test. (D) (p)ppGpp, RNAP site 1, and not site 2, promote the cipro-induced SOS response. 16 h growth in MAC cipro (log phase). (E) (p)ppGpp and RNAP sites 1 and 2 are required for cipro-induction of stringent response transcription: site 2, presumably, because it is required for the transcriptional program, and site 1 because of its defect in SOS (D, above), both of which are necessary for cipro-induction of the stringent response-on subpopulation. 16 h growth in MAC cipro (log phase). (F) RNAP sites 1 and 2 are required for cipro-induction of the σ^S^-active gambler-cell subpopulation. 16 h growth in MAC cipro (log phase). (D-F) Left, representative flow-cytometry histograms. Right, means ± SEM, ≥ 3 experiments. *p<0.01 (D), *p<0.001 (B, E, F), n.s. not significant, one-way ANOVA with Tukey’s post-hoc test. (G) Model for (p)ppGpp and RNAP site 1 promotion of SOS induction by stabilizing backtracked RNAP. (i) DSB in DNA being transcribed. (ii) Site 1 and (p)ppGpp stabilize RNAP backtracked at a transcription error; backtracked RNAP stalls RecBCD degradation, switching it to ssDNA end creation and (iii) RecA loading, which promotes both repair and the SOS response. This model is based on findings of (Sivaramakrishnan et al., 2017), and ours regarding (p)ppGpp- and RNAP site 1-dependence of cipro-induced SOS induction. This model implies that a large fraction of cipro-induced DSB ends is resected into actively transcribed genes where they are paused by RNAP, contributing to SOS and repair.

Next, RNAP site 2 (transcriptional program) played no role in surviving MAC cipro (8.5 ng/ml, Figure 5C), whereas site 1 (backtracking) promoted survival, discussed below. Similarly, site 2 (transcription) had no significant effect on SOS induction by cipro (Figure 5D); but site 2 promoted cipro-induced stringent-response transcription of the reporter (Figure 5E), and induction of σ^S^ activity (Figure 5F). These data agree with the roles of DksA (transcription) summarized Figure 4E; acting solely after ROS, the transcriptional program upregulates the sRNAs that activate the σ^S^ response and MBR.

Finally, in agreement with the role of site 1 (backtracking) in survival of MAC cipro (Figure 5C), site 1 also made a modest but significant contribution to cipro induction of SOS (Figure 5D, as did (p)ppGpp, shown with the (p)ppGpp-null, Δ*relA ΔspoT* double mutant (Figure 5D). The data support a model in which backtracked RNAP promotes SOS induction and survival, which, presumably, occurs by DSB repair. Backtracked RNAP has been shown to block DNA degradation from DSBs and promotes DSB repair (Sivaramakrishnan et al., 2017). The switch to DSB repair requires loading of RecA onto DNA (Churchill et al., 1999), and the resulting RecA-DNA complex is the inducer of the SOS response (Sassanfar and Roberts, 1990) (Figure 5G, Discussion). Increased RecA loading promoted by backtracked RNAP could explain the requirement for (p)ppGpp in cipro-induced SOS activity (Figure 4A). The data support a model (Figure 5G) in which (p)ppGpp in RNAP site 1 stabilize backtracked RNAP, blocking degradation by RecBCD and promoting RecA loading, which both activates SOS and is required for DSB repair, and so, also, for MBR.

## DISCUSSION

The data presented identify the stringent starvation stress response and RNA polymerase as critical regulators of the cipro-induced pathway to transient differentiation of a mutable gambler-cell subpopulation, which generates antibiotic-resistant and cross-resistant mutant bacteria (Pribis et al., 2019). The stringent response transcriptional program is induced by ROS (Figures 2A and 2B), then upregulates two sRNAs (Figure 2C) that allow activation of σ^S^ (Figures 1B and 1C) (Pribis et al., 2019). The ROS-high cell subpopulation emerges from a whole population of SOS-induced cells (Pribis et al., 2019), and then becomes stringent-active (Figure 3A), then σ^S^-active gambler cells (Figures 3B and 4E). Gamblers “risk” mutagenesis while other cells maintain genome stability (Pribis et al., 2019), a potential “bet-hedging” strategy (Pribis et al., 2019; Torkelson et al., 1997; Veening et al., 2008). The steps in the genesis of gamblers are attractive targets for drugs to slow evolution of resistance (Pribis et al., 2022; Waylen et al., 2020), and those steps identified here expand options for finding such drugs.

The data also revealed that stringent-response activator (p)ppGpp and RNAP promote DNA-damage signaling in the cipro-induced SOS response (Figures 4A, C and 5). Promotion of SOS and mutagenesis by RNAP site 1 (Figures 5B ands 5D) implies that RNAP, and not some previously unknown other target, underlies the (p)ppGpp role(s) in SOS and mutagenesis. In Figure 5G, we propose that (p)ppGpp and RNAP facilitate the switch from RecBCD degradation of DSB ends to loading RecA onto single-stranded (ss)DNA, directly, by stabilizing backtracked RNAP at the site of DNA breakage (Figure 5Gii). Previous work showed that backtracked RNAP pauses RecBCD DSB-end destruction, which, like pausing at Chi sites, shifts RecBCD to DSB-repair mode, ending DNA destruction and inducing repair of the DSB (Sivaramakrishnan et al., 2017). RecBCD pauses, then exposes ssDNA onto which it loads RecA (Churchill et al., 1999), which both creates the SOS-inducing signal—a RecA-DNA nucleoprotein complex (Sassanfar and Roberts, 1990)—and begins DSB repair (Figure 5Giii) (Sivaramakrishnan et al., 2017). We hypothesize that (p)ppGpp and RNAP site 1 (backtracking) upregulate cipro-induced SOS activity, DSB repair and mutagenesis by stabilizing backtracked RNAP causing a direct physical blockage to, and pausing of, RecBCD (Figure 5G). (p)ppGpp also promotes nucleotide excision repair (Kamarthapu et al., 2016), a process unrelated to the homology-directed repair that mends DSBs, studied by (Sivaramakrishnan et al., 2017). This model (Figure 1G) implies that starvation, as sensed by the stringent response, favors preservation of broken DNA by repair—a potentially beneficial counter to the findings of (Sivaramakrishnan et al., 2017), in which rapid growth in rich medium favored transcription fidelity at the expense of DSB repair and chromosome loss. Rapidly growing *E. coli*, with up to 8 chromosomes per cell, can afford the loss of the broken chromosome, and instead protect the proteome that will be inherited by all 7 or 8 future cells (Sivaramakrishnan et al., 2017). Few chromosomes are present in starving cells (Akerlund et al., 1995) and their preservation during starvation appears to be favored.

The data also suggest that transcription is likely to shape the genomic landscape of cipro-induced mutagenesis. Cipro inhibits type-II topoisomerases, which cleave DNA to relieve supercoils during both replication and transcription (Anderson and Osheroff, 2001). The role of RNAP site 1 (backtracking) in SOS induction (Figure 5D) and survival, presumably reflecting DSB repair (Figure 5C), suggests that cipro-induced DNA breakage often occurs in sites occupied by RNAP, *i*.*e*, within active genes. This model (Figure 5G) implies that cipro-induced MBR is likely to be biased to genomic regions with actively transcribed genes. Increased mutagenesis in and near active genes could speed evolution of genes in use in a given environment, a potential means of accelerating adaptation.

## ACKNOWLEDGMENTS

We thank Devon Fitzgerald for comments on the manuscript. Supported by National Institutes of Health (NIH) grants R35-GM122598 and R01-CA250905 (SMR), R01-GM106373 (PJH), and NIH Directors Pioneer Awards DP1-AI52073 (CH) and DP1-AG072751 (SMR), the BCM Cytometry and Cell Sorting Core through CPRIT Core Facility Support Award (CPRIT-RP180672) and NIH (CA125123 and RR024574) aided by JM Sederstrom.

## Key Resources Table

| REAGENT or RESOURCE | SOURCE | IDENTIFIER |
| --- | --- | --- |
| **Chemicals** |  |  |
| ciprofloxacin | MP Biomedicals | Cat# 199020 |
| rifampicin | Research Products International | Cat# 13292-46-1 |
| kanamycin | Sigma-Aldrich | Cat# K1876 |
| ampicillin | Sigma-Aldrich | Cat# A9518 |
| thiourea | Sigma-Aldrich | Cat# T8656 |
| formaldehyde | Sigma-Aldrich | Cat# 252549 |
| dihydrorhodamine | Life Technologies | Cat# D632 |
| 2-Nitrophenyl β-D-galactopyranoside | Sigma-Aldrich | Cat# N1127 |
| Experimental Models | *Escherichia coli* strains Table S1 | N/A |

## Methods Details

### Bacterial Strains, Media, and Growth

*E. coli* strains used, and their origins are given in Table S1. Specific strains used in each main figure are listed in the following section. Assays for rifampicin-resistant (RifR) and ampicillin-resistant (AmpR) mutants were performed in one of two wild-type (WT) *E. coli* strains, and their isogenic derivatives. Bacteria were grown in LBH rich medium at 37°C with aeration, and additives where indicated at the following concentrations: ciprofloxacin (cipro, 1–8.5 ng/mL, Table S2), thiourea (100mM), ampicillin (100μg/ml), chloramphenicol (25μg/ml), kanamycin (50μg/ml), tetracycline (10μg/ml), rifampicin (110μg/ml), and sodium citrate (20mM). The minimum antibiotic concentration, or MAC, is the concentration at which 10% of treated cells remain viable, compared with cultures with no antibiotic (Lorian and De Freitas, 1979), and was determined for each strain experimentally, as previously (Pribis et al., 2019).

### Strains Used in Figures 1-5

Fig. 1: (B) SMR24096, SMR24134, SMR24441, SMR24447. (C) SG30013, SMR26995, SMR27001. (D) MG1655, SMR20469, SMR20471, SMR27086, SMR5223, SMR23927, SMR23919, SMR27084. Fig. 2: (A) SMR24273, SMR27003, SMR27009. (B) SMR24273, SMR27009. (C) CH2046, SMR27013, SMR27019, PM1450, SMR27021, SMR27027. (D) SMR24273, SMR27009, SMR27033, SMR27035, SMR27088. Fig. 3: (A) SMR27039. (B) SMR27011. Fig. 4: (A) SMR24100, SMR24156, SMR27043. (B) SMR24100, SMR24156, SMR27041. (C) MG1655, SMR20471, SMR24426. (D) MG1655, SMR20471. (F) SMR24273, SMR27009, SMR27029, SMR27031. Fig. 5: (B) MG1655, CH6670, CH6669. (C) MG1655, CH6670, CH6669. (D) SMR24100, SMR27072, SMR27074, SMR27043, SMR24156. (E) SMR24273, SMR27076, SMR27078, SMR27009. (F) SMR24096, SMR27080, SMR27082, SMR24134.

### Assays for Ciprofloxacin-induced Mutagenesis

Assays for rifampicin-resistant (RifR) and ampicillin-resistant (AmpR) mutants were performed with cipro at MAC in the wild-type (WT) *E. coli* strains MG1655 and SMR5223, and their isogenic derivates, as previously (Pribis et al., 2019). Ten to sixty aliquots of log-phase cultures were diluted 1:3 and dispensed into 14-mL tubes with and without MAC cipro, and then grown at 37°C shaking for the mutagenesis assays for 24 h (RifR) or 48 h (AmpR) then assayed for mutant and total CFU per (Pribis et al., 2019). Mutation rates were estimated with the MSS-MLE algorithm using the FALCOR calculator (Hall et al., 2009).

### Flow Cytometry for SOS and σ^S^ Activity

Flow cytometric assays for SOS- and σ^S^-response-regulated promoter activity were as described (Pribis et al., 2019). Quantification of cells that have induced their SOS or σ^S^ response, and how much they have, was achieved using engineered chromosomal fluorescence reporter genes and flow cytometry, per (Nehring et al., 2016; Pennington and Rosenberg, 2007) for SOS, and per (Al Mamun et al., 2012; Pribis et al., 2019) for σ^S^-response activation. We used the *yiaG-yfp* σ^S^-response reporter and the *Δattλ*::P_*sulA*_*mCherry* SOS reporter (Nehring et al., 2016) modified from (Pennington and Rosenberg, 2007) in strains grown under fluctuation-test conditions as described for Assays for Ciprofloxacin-induced Mutagenesis, with or without cipro, at indicated concentration(s), and cells were harvested in late log phase or stationary phase. For quantification, flow cytometry “gates” were calibrated, for SOS, using the negative-control SOS-off *lexA*(Ind^-^), and SOS-response proficient cells (Pennington and Rosenberg, 2007) as the dividing place between peaks of the distribution of SOS-proficient cells at which most cells diverge from the spontaneously SOS-induced fluorescent cell subpopulation, usually at between 0.5% and 1% of cells cultured in LBH broth. With this gate, ∼10^−4^ of SOS-non-inducible *lexA*Ind^-^ cells cross the gate, scoring as “SOS-positive” (Pennington and Rosenberg, 2007). For the σ^S^ response, gates were set to the point at which fewer than 0.5% of cells with cipro but without the reporter gene were positive. At this gate, fewer than 10^−3^ of Δ*rpoS* cells, which are σ^S^-response deficient, cross the gate and would be scored as positive. For all, the percent of the population that scored as positive is reported.

### Stringent Response Fluorescence Reporter Construction and Flow Cytometry

mCherry FRT*cat*FRT was amplified from strain SMR21793 and moved to nucleotide position *zfe2512*.*7* in the *E. coli* genome, between the divergently transcribed *mntH* and *nupC* genes by short-homology phage lambda Red-mediated recombineering using primers P1 and P2 and verified with Primers P3 and P4 (Table S3). For the stringent response, gates for stringent response-active cells were set to the point at which fewer than 0.5% of cells with cipro but without the reporter gene were positive. At this gate, fewer than 10^−3^ of Δ*relA* Δ*spoT* negative control (p)ppGpp^0^ cells, which are stringent-response deficient, cross the gate and would be scored as positive.

### Single-Cell Detection of Intracellular ROS by Flow Cytometry

Cells were grown in the absence or presence of cipro at its MAC (8.5ng/mL) as for Assays for Ciprofloxacin-induced Mutagenesis. The ROS measurement protocol was adapted from Pribis et al (Pribis et al., 2019). Cells were harvested serially from cultures for ROS detection using dihyrdorhodamine (DHR, Life Technologies), or using transcriptional fusions of the *oxyR* promoters to GFP. For ROS detection using P_*oxyR*_*gfp*, cells containing stringent-response reporter P_*rmf*_*mCherry* and plasmids carrying the P_*oxyR*_*gfp* transcriptional fusions, or a promoterless *gfp* parental plasmid Pvector-*gfp*, were maintained with kanamycin, and used to detect both GFP and red fluorescence. Single color controls were also collected at time points for spectral compensation. For the P_*oxyR*_*gfp*, gates were drawn so that the promoterless-*gfp* vector Pvector-*gfp* had 0.5% GFP-positive cells.

### Beta-galactosidase Assays

Cells were grown as for Assays for Ciprofloxacin-induced Mutagenesis to equivalent ODs and frozen at -20ºC until assays were carried out. Determination of the β-galactosidase activity of the P_*dsrA*_*-lacZ*, P_*arcZ*_*-lacZ, rpoS-lacZ* fusions was accomplished using the standard assay described by JH Miller, as previously (Gibson et al., 2010; Miller, 1992; Pribis et al., 2019).

### Statistics

For comparisons of two groups, a two-tailed Students t-test was used. For comparisons of 3 or more groups, ANOVA with Tukey post-hoc test was used. Statistics were performed using GraphPad PRISM.

## REFERENCES

Akerlund, T., Nordstrom, K., and Bernander, R. (1995). Analysis of cell size and DNA content in exponentially growing and stationary-phase batch cultures of Escherichia coli. J Bacteriol 177, 6791–6797.

Al Mamun, A.A., Lombardo, M.J., Shee, C., Lisewski, A.M., Gonzalez, C., Lin, D., Nehring, R.B., Saint-Ruf, C., Gibson, J.L., Frisch, R.L., et al. (2012). Identity and function of a large gene network underlying mutagenic repair of DNA breaks. Science 338, 1344–1348.

Anderson, V.E., and Osheroff, N. (2001). Type II topoisomerases as targets for quinolone antibacterials: turning Dr. Jekyll into Mr. Hyde. Curr Pharm Des 7, 337–353.

Ardal, C., Balasegaram, M., Laxminarayan, R., McAdams, D., Outterson, K., Rex, J.H., and Sumpradit, N. (2020). Antibiotic development - economic, regulatory and societal challenges. Nat Rev Microbiol 18, 267–274.

Battesti, A., Majdalani, N., and Gottesman, S. (2015). Stress sigma factor RpoS degradation and translation are sensitive to the state of central metabolism. Proc. Natl Acad Sci. USA 112, 5159–5164.

Blair, J.M., Webber, M.A., Baylay, A.J., Ogbolu, D.O., and Piddock, L.J. (2015). Molecular mechanisms of antibiotic resistance. Nat Rev Microbiol 13, 42–51.

Blazquez, J., Rodriguez-Beltran, J., and Matic, I. (2018). Antibiotic-induced genetic variation: how it arises and how it can be prevented. Annu Rev Microbiol 72, 209–230.

Churchill, J.J., Anderson, D.G., and Kowalczykowski, S.C. (1999). The RecBC enzyme loads RecA protein onto ssDNA asymmetrically and independently of chi, resulting in constitutive recombination activation. Genes Dev 13, 901–911.

Cirz, R.T., Chin, J.K., Andes, D.R., de Crecy-Lagard, V., Craig, W.A., and Romesberg, F.E. (2005). Inhibition of mutation and combating the evolution of antibiotic resistance. PLoS Biol 3, e176.

Cullen, M.E., Wyke, A.W., Kuroda, R., and Fisher, L.M. (1989). Cloning and characterization of a DNA gyrase A gene from Escherichia coli that confers clinical resistance to 4-quinolones. Antimicrob Agents Chemother 33, 886–894.

Davies, J., and Davies, D. (2010). Origins and evolution of antibiotic resistance. Microbiol Mol Biol Rev 74, 417–433.

Drlica, K. (1999). Mechanism of fluoroquinolone action. Curr Opin Microbiol 2, 504–508.

Gibson, J.L., Lombardo, M.J., Thornton, P.C., Hu, K.H., Galhardo, R.S., Beadle, B., Habib, A., Magner, D.B., Frost, L.S., Herman, C., et al. (2010). The sigma(E) stress response is required for stress-induced mutation and amplification in Escherichia coli. Mol Microbiol 77, 415–430.

Girard, M.E., Gopalkrishnan, S., Grace, E.D., Halliday, J.A., Gourse, R.L., and Herman, C. (2018). DksA and ppGpp regulate the sigma(S) stress response by activating promoters for the small RNA DsrA and the anti-Adapter protein IraP. J Bacteriol 200.

Govindaraj Vaithinathan, A., and Vanitha, A. (2018). WHO global priority pathogens list on antibiotic resistance: an urgent need for action to integrate One Health data. Perspect Public Health 138, 87–88.

Gutierrez, A., Laureti, L., Crussard, S., Abida, H., Rodriguez-Rojas, A., Blazquez, J., Baharoglu, Z., Mazel, D., Darfeuille, F., Vogel, J., et al. (2013). beta-Lactam antibiotics promote bacterial mutagenesis via an RpoS-mediated reduction in replication fidelity. Nat Commun 4, 1610.

Hall, B.M., Ma, C.X., Liang, P., and Singh, K.K. (2009). Fluctuation analysis CalculatOR: a web tool for the determination of mutation rate using Luria-Delbruck fluctuation analysis. Bioinformatics 25, 1564–1565.

Howard, B.M., Pinney, R.J., and Smith, J.T. (1993). Function of the SOS process in repair of DNA damage induced by modern 4-quinolones. J Pharm Pharmacol 45, 658–662.

Izutsu, K., Wada, A., and Wada, C. (2001). Expression of ribosome modulation factor (RMF) in Escherichia coli requires ppGpp. Genes Cells 6, 665–676.

Kamarthapu, V., Epshtein, V., Benjamin, B., Proshkin, S., Mironov, A., Cashel, M., and Nudler, E. (2016). ppGpp couples transcription to DNA repair in E. coli. Science 352, 993–996.

Kohanski, M.A., DePristo, M.A., and Collins, J.J. (2010). Sublethal antibiotic treatment leads to multidrug resistance via radical-induced mutagenesis. Mol Cell 37, 311–320.

Komissarova, N., and Kashlev, M. (1997). Transcriptional arrest: Escherichia coli RNA polymerase translocates backward, leaving the 3’ end of the RNA intact and extruded. Proc Natl Acad Sci U S A 94, 1755–1760.

Lombardo, M.J., Aponyi, I., and Rosenberg, S.M. (2004). General stress response regulator RpoS in adaptive mutation and amplification in Escherichia coli. Genetics 166, 669–680.

Lorian, V., and De Freitas, C.C. (1979). Minimal antibiotic concentrations of aminoglycosides and beta-lactam antibiotics for some gram-negative bacilli and gram-positive cocci. J Infect Dis 139, 599–603.

Martin, J.K., 2nd, Sheehan, J.P., Bratton, B.P., Moore, G.M., Mateus, A., Li, S.H., Kim, H., Rabinowitz, J.D., Typas, A., Savitski, M.M., et al. (2020). A dual-mechanism antibiotic kills gram-negative bacteria and avoids drug resistance. Cell 181, 1518–1532 e1514.

McPartland, A., Green, L., and Echols, H. (1980). Control of recA gene RNA in E. coli: regulatory and signal genes. Cell 20, 731–737.

Mehrad, B., Clark, N.M., Zhanel, G.G., and Lynch, J.P., 3rd (2015). Antimicrobial resistance in hospital-acquired gram-negative bacterial infections. Chest 147, 1413–1421.

Miller, J.H. (1992). A short course in bacterial genetics : a laboratory manual and handbook for Escherichia coli and related bacteria (Plainview, N.Y.: Cold Spring Harbor Laboratory Press).

Minnick, P.J. (2019). ppGpp orchestrates mutagenesis by mediating transcription-DNA repair conflict. (Ph.D. thesis, Department of Biochemistry and Molecular Biology, Baylor College of Medicine).

Murray, C.J.L., Ikuta, K.S., Sharara, F., Swetschinski, L., Robles Aguilar, G., Gray, A., Han, C., Bisignano, C., Rao, P., Wool, E., et al. (2022). Global burden of bacterial antimicrobial resistance in 2019: a systematic analysis. The Lancet.

Nehring, R.B., Gu, F., Lin, H.Y., Gibson, J.L., Blythe, M.J., Wilson, R., Bravo Nunez, M.A., Hastings, P.J., Louis, E.J., Frisch, R.L., et al. (2016). An ultra-dense library resource for rapid deconvolution of mutations that cause phenotypes in Escherichia coli. Nucleic Acids Res 44, e41.

Novogrodsky, A., Ravid, A., Rubin, A.L., and Stenzel, K.H. (1982). Hydroxyl radical scavengers inhibit lymphocyte mitogenesis. Proc. Natl Acad Sci. USA 79, 1171–1174.

Oethinger, M., Podglajen, I., Kern, W.V., and Levy, S.B. (1998). Overexpression of the marA or soxS regulatory gene in clinical topoisomerase mutants of Escherichia coli. Antimicrob Agents Chemother 42, 2089–2094.

Oram, M., and Fisher, L.M. (1991). 4-Quinolone resistance mutations in the DNA gyrase of Escherichia coli clinical isolates identified by using the polymerase chain reaction. Antimicrob Agents Chemother 35, 387–389.

Pennington, J.M., and Rosenberg, S.M. (2007). Spontaneous DNA breakage in single living Escherichia coli cells. Nat Genet 39, 797–802.

Ponder, R.G., Fonville, N.C., and Rosenberg, S.M. (2005). A switch from high-fidelity to error-prone DNA double-strand break repair underlies stress-induced mutation. Mol Cell 19, 791–804.

Pribis, J.P., Garcia-Villada, L., Zhai, Y., Lewin-Epstein, O., Wang, A.Z., Liu, J., Xia, J., Mei, Q., Fitzgerald, D.M., Bos, J., et al. (2019). Gamblers: an antibiotic-induced evolvable cell subpopulation differentiated by reactive-oxygen-induced general stress response. Mol Cell 74, 785–800 e787.

Pribis, J.P., Zhai, Y., Hastings, P.J., and Rosenberg, S.M. (2022). Stress-induced mutagenesis, gambler cells, and stealth targeting antibiotic-induced evolution. mBio, e0107422.

Rosenberg, S.M., and Queitsch, C. (2014). Medicine. Combating evolution to fight disease. Science 343, 1088–1089.

Ross, W., Sanchez-Vazquez, P., Chen, A.Y., Lee, J.H., Burgos, H.L., and Gourse, R.L. (2016). ppGpp binding to a site at the RNAP-DksA interface accounts for its dramatic effects on transcription initiation during the stringent response. Mol Cell 62, 811–823.

Ross, W., Vrentas, C.E., Sanchez-Vazquez, P., Gaal, T., and Gourse, R.L. (2013). The magic spot: a ppGpp binding site on E. coli RNA polymerase responsible for regulation of transcription initiation. Mol Cell 50, 420–429.

Sanchez-Vazquez, P., Dewey, C.N., Kitten, N., Ross, W., and Gourse, R.L. (2019). Genome-wide effects on Escherichia coli transcription from ppGpp binding to its two sites on RNA polymerase. Proc Natl Acad Sci U S A 116, 8310–8319.

Sassanfar, M., and Roberts, J.W. (1990). Nature of the SOS-inducing signal in Escherichia coli. The involvement of DNA replication. J Mol Biol 212, 79–96.

Shee, C., Gibson, J.L., Darrow, M.C., Gonzalez, C., and Rosenberg, S.M. (2011). Impact of a stress-inducible switch to mutagenic repair of DNA breaks on mutation in Escherichia coli. Proc Natl Acad Sci U S A 108, 13659–13664.

Shee, C., Gibson, J.L., and Rosenberg, S.M. (2012). Two mechanisms produce mutation hotspots at DNA breaks in Escherichia coli. Cell Rep 2, 714–721.

Sivaramakrishnan, P., Sepulveda, L.A., Halliday, J.A., Liu, J., Nunez, M.A.B., Golding, I., Rosenberg, S.M., and Herman, C. (2017). The transcription fidelity factor GreA impedes DNA break repair. Nature 550, 214–218.

Torkelson, J., Harris, R.S., Lombardo, M.J., Nagendran, J., Thulin, C., and Rosenberg, S.M. (1997). Genome-wide hypermutation in a subpopulation of stationary-phase cells underlies recombination-dependent adaptive mutation. EMBO J 16, 3303–3311.

Touati, D., Jacques, M., Tardat, B., Bouchard, L., and Despied, S. (1995). Lethal oxidative damage and mutagenesis are generated by iron in delta fur mutants of Escherichia coli: protective role of superoxide dismutase. J Bacteriol 177, 2305–2314.

Ubukata, K., Itoh-Yamashita, N., and Konno, M. (1989). Cloning and expression of the norA gene for fluoroquinolone resistance in Staphylococcus aureus. Antimicrob Agents Chemother 33, 1535–1539.

Veening, J.W., Smits, W.K., and Kuipers, O.P. (2008). Bistability, epigenetics, and bet-hedging in bacteria. Annu Rev Microbiol 62, 193–210.

Wang, J.C. (1998). Moving one DNA double helix through another by a type II DNA topoisomerase: the story of a simple molecular machine. Q Rev Biophys 31, 107–144.

Wang, Z., Koirala, B., Hernandez, Y., Zimmerman, M., and Brady, S.F. (2022). Bioinformatic prospecting and synthesis of a bifunctional lipopeptide antibiotic that evades resistance. Science 376, 991–996.

Waylen, L.N., Nim, H.T., Martelotto, L.G., and Ramialison, M. (2020). From whole-mount to single-cell spatial assessment of gene expression in 3D. Commun Biol 3, 602.

Werner, N.L., Hecker, M.T., Sethi, A.K., and Donskey, C.J. (2011). Unnecessary use of fluoroquinolone antibiotics in hospitalized patients. BMC Infect Dis 11, 187.

Zaslaver, A., Bren, A., Ronen, M., Itzkovitz, S., Kikoin, I., Shavit, S., Liebermeister, W., Surette, M.G., and Alon, U. (2006). A comprehensive library of fluorescent transcriptional reporters for Escherichia coli. Nat Methods 3, 623–628.

Zhang, Q., Lambert, G., Liao, D., Kim, H., Robin, K., Tung, C.K., Pourmand, N., and Austin, R.H. (2011). Acceleration of emergence of bacterial antibiotic resistance in connected microenvironments. Science 333, 1764–1767.

